# Sensitive period for song learning outlasts parent-offspring cohesion in wild zebra finches

**DOI:** 10.64898/2026.10.28.762438

**Authors:** Noëlle Tschirren, Hugo Loning, Lysanne Snijders, Simon Griffith, Marc Naguib, Chris Tyson

## Abstract

Communication is central to social behaviour, with vocalisations playing a crucial role in many species. In some species, vocalisations are learned, with songbirds being key to understanding vocal learning. The Australian zebra finch (*Taeniopygia castanotis*) is the most studied model organism for vocal learning. In captivity, juveniles are typically housed with their father, where males learn their father’s song and females develop a preference for that song. In nature, however, the social structure of zebra finches is more dynamic across space and time. To identify the natural social conditions during song learning, we followed zebra finch families across their early life in the wild using automated radio- tracking. The distance between juveniles and their parents increased post-fledging and became similar to the distance between juveniles and unrelated adults by the second half of the song learning period. These findings indicate a highly dynamic social environment during the song learning period and suggest that juvenile males could learn from many tutors and that females may develop broader song preferences. Thus, studies based on one-tutor paradigms should be interpreted carefully with respect to functional implications and highlight that future vocal learning studies could better reflect the social conditions under which song develops.

---

Across taxa, most animals use species-specific vocalisations in order to communicate with each other (Bradbury and Vehrencamp, 2011). In most species, these vocalisations develop without major social input. Contrastingly, in a few taxa, including humans, cetaceans, bats, and three orders of birds, vocalisations are learned primarily early in life, and therefore shaped by the early social environment (Janik, 2014; Nowicki and Searcy, 2014; Wilbrecht and Nottebohm, 2003). If juveniles are not exposed to species-specific vocalisations during this period, their own vocalisations deviate strongly from conspecifics who had the opportunity to learn (Brenowitz and Beecher, 2005; Marler and Sherman, 1985; Prat et al., 2015; Slater, 1989), which can result in negative fitness effects (Crates et al., 2021). Songbirds are the most extensively studied vocal learners and serve as a key model system for understanding the neurobiology and development of social learning, with many parallels drawn to human speech acquisition (Hyland Bruno et al., 2021; Jarvis, 2019). While some species are open-ended learners, defined by having the ability to continue to add or modify song elements throughout life, others are closed-ended learners, where song learning is constrained to a limited developmental period early in life (Beecher and Brenowitz, 2005; Brainard and Doupe, 2002; Hultsch and Todt, 2004).

In species that learn their vocalisations from conspecifics early in life, the social environment and auditory landscape in which juveniles develop determine which potential tutors are encountered and, consequently, which vocalisations can be learned. The social environment of juveniles can change substantially during the transition to independence, and tutor exposure is likely to change over the course of development. As such, neuronal and behavioural mechanisms for song learning have evolved under diverse social conditions. Juveniles from territorial songbirds, for instance, are initially isolated from neighbours, but after fledging may move around substantially during the song learning period, where they may seek diverse input for learning (Templeton et al., 2012). In some species, only few renditions need to be heard (Hultsch and Todt, 2004), which can be adaptive in territorial species where males sing much less during the period of raising offspring compared to early in the season (Amrhein et al., 2002; Catchpole, 2000). In such cases, an extended window for vocal learning can be beneficial (Liu and Kroodsma, 2006; Mennill et al., 2018; Wheelwright et al., 2008), enabling a juvenile to learn the song of close neighbours after dispersal, or even extending to several years, with song repertoires increasing with age (Kiefer et al., 2010, 2009).

Across all non-human animals studied for vocal learning, and specifically among songbirds, the zebra finch (*Taeniopygia castanotis*) has been the prime model organism across laboratories worldwide (Griffith et al., 2021; Hauber et al., 2021; Hyland Bruno et al., 2021; Zann, 1996). As a result, zebra finch vocal learning is one of the best-studied learned behaviours in non-human animals, which has generated thousands of publications across many decades of research (e.g. Griffith and Buchanan, 2010; Hauber et al., 2021; Morris, 1954). This research has provided key insights into the developmental and neuronal mechanisms underlying song perception and production as well as social aspects affecting song learning by males and song preference learning by females (Hauber et al., 2021; Hyland Bruno et al., 2021). A key advantage of this model is that males sing one individually distinct song motif that they acquire through exposure to adult males during a sensitive developmental period early in life. In zebra finches, the sensitive period of song acquisition typically occurs from 25-65 days post-hatching (Doupe and Kuhl, 1999; Eales, 1985; Gobes et al., 2019; Immelmann, 1969). Once sexual maturity is reached, the learned song motive becomes crystallised and thereafter remains largely unchanged (Böhner, 1983; Immelmann, 1969; Tchernichovski et al., 2001). Much of the existing work on the mechanisms of song learning in zebra finches comes from laboratory studies, which provide highly stable and restricted social environments (Hauber et al., 2021). In captive settings, juveniles typically are housed with a single tutor, often the father, and juvenile males are generally found to learn their fathers’ song (Böhner, 1983; Clayton, 1987; Mann and Slater, 1995, 1994; Roper and Zann, 2006), while juvenile females develop preferences for songs like their father’s (Honarmand et al., 2015; Riebel, 2009, 2000). However, song copying is not exact, with juvenile males introducing individual-specific variation into their song (Böhner, 1983). Additionally, when reared in larger aviaries with several breeding pairs, juvenile males can learn song from multiple males (Eales, 1985; Williams, 1990). Together, these findings imply that zebra finches are able and motivated to learn from more than a single tutor, and that song-learning paradigms with a single tutor may not be the most ecologically relevant.

In free-living populations, the social structure, which determines the availability of tutors for song learning, is far more variable than in laboratory settings. Zebra finches typically move around in pairs or small mixed-sex groups, but aggregate during socialising and foraging and breed colonially, resulting in a dynamic fission-fusion social system (Brandl et al., 2021; Hagedoorn et al., 2026; McCowan et al., 2015; Tyson et al., 2024). While some zebra finches can breed far from colonies and may be isolated, birds in colonies usually assemble at social hotspots that likely facilitate social information exchange (Loning et al., 2023). Such fluid social environments expose juveniles to many individuals and a wide range of vocalisations, substantially increasing the potential pool of tutors beyond the father. Thus, understanding the social structure during the sensitive period of development is critical for understanding the factors shaping the ontogeny of vocal behaviour and helps place laboratory findings within a more realistic ecological context.

Here, we thus tested how the social environment of wild juvenile zebra finches in the Australian outback changed over the course of ontogeny. We tracked how juvenile spatial proximity to parents, siblings, and other members in the colony changed over post-fledging development, and therefore across the sensitive period of song acquisition. Using an automated radio-tracking system, we monitored the movements of 29 families from juvenile fledging until after the sensitive period of song acquisition. We expected that after fledging, juveniles would initially stay in close spatial proximity to their parents, moving throughout the environment together, as during this time juveniles are still provisioned by their parents. We further predicted that parent-offspring distance would gradually increase throughout the sensitive period of song learning, rather than only after its end, as juveniles are able to forage independently before this time and parents may initiate another breeding attempt within a few weeks of fledging their offspring (Zann 1996). A gradual decline in family cohesion, together with gradually being closer to other birds, would increase opportunities for juveniles to encounter a diverse set of adults and other juveniles and, consequently, to acquire song elements from individuals other than their father. Additionally, we assessed whether juvenile males and females differ in their proximity to parents, and whether juveniles differed in their distance to their mother versus their father, given that the benefits of remaining close may vary by sex.

## Methods

### Field procedures

Fieldwork took place at the Fowlers Gap research station in New South Wales, Australia (31°05ʹ25ʺS 141°42ʹ30ʺE), between September and December 2024, coinciding with the peak breeding activity of zebra finches in this area. At the local field site ’Gap Hills’, we utilised nest boxes installed previously (Griffith et al., 2008). Nest boxes were checked every three days, which allowed for a precise identification of the onset of new breeding activity as females lay one egg on each consecutive day, and the usual clutch size averages 5 eggs (Griffith et al., 2008). In wild zebra finches, nestlings from one brood hatch synchronously on one or across two consecutive days (Mainwaring et al., 2010). As such, the first hatching day was marked as day 0 for that brood (Fig. 1). Nestlings are regularly visited by both parents for feeding, which allowed us to capture parents visiting their offspring at the nest from day 7 on. Birds were weighed with a 20 g Pesola and equipped with a uniquely identifiable bird band and a solar-powered radio tag attached via a leg harness (Cellular Tracking Technologies LifeTag, NJ, USA). The first parent was caught between day 7 and day 9, the second parent was captured with at least one day in between, between day 9 and 11 (Fig. 1). This allowed the first parent to become accustomed to the tag before equipping the second parent, ensuring uninterrupted feeding for the nestlings. When the second parent was too cautious to enter the nest with the trap installed, only the first parent was tagged. Once one or both parents were tagged, the nest was left undisturbed until day 16, when nestlings were weighed, and a breast feather sample was taken to determine sex (Curtin University DNA Sexing). While preferably all nestlings were equipped with a tag, too many tags in one nest risked antenna entanglement, so we equipped only two nestlings of each brood with a tag. Nestlings generally fledge 18-20 days after hatching, after which both parents continue to feed their offspring outside of the nest (Zann, 1996). To increase the sample size of tagged juveniles, we tracked fledged tagged juveniles with handheld locators (Cellular Tracking Technologies Locator, NJ, USA) and caught their untagged siblings using mist nets to equip them with tags as well. This resulted in a mean of 2.8 juveniles tagged per nest (min = 1, max = 6).

**Figure 1.**
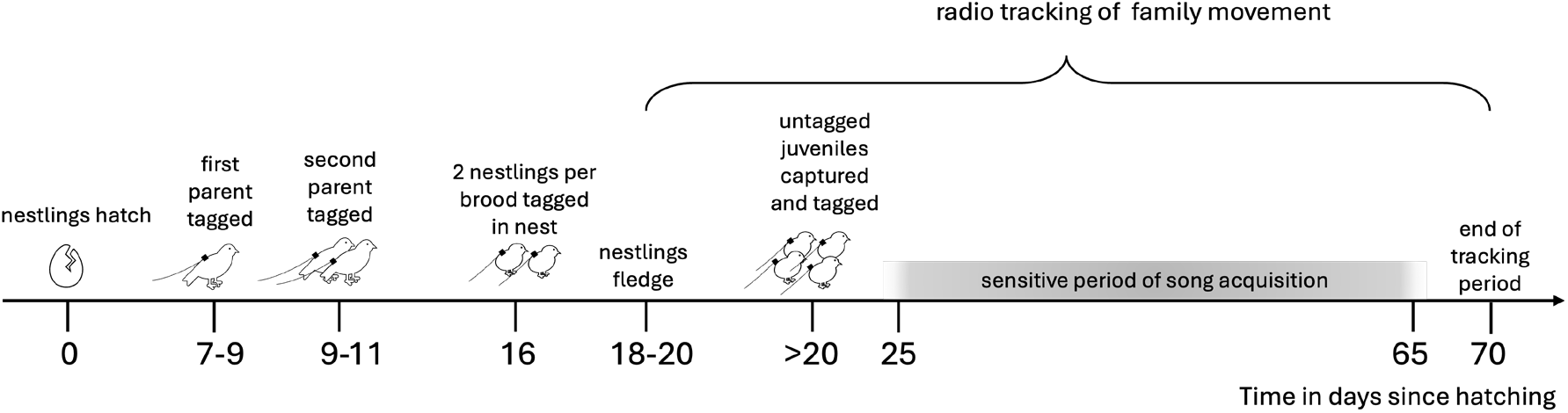
Schematic timeline of radio tracking in relation to juvenile development. The sensitive period of song acquisition as identified in laboratory studies is between days 25 and 65 post-hatching.

### Automated radio tracking

The radio tags weighed an average of 0.53 g ± 0.03 g, which was below 5% of the body weight measured across adult birds (12.4 g ± 0.9 g). Tags were programmed to signal every 5 s, which was picked up by a grid of 109 radio receiver nodes (Cellular Tracking Technologies Node v. 2, NJ, USA), spaced ≤ 150 m apart, covering an area of approximately 1.5 km^2^ (Fig. 2). Nodes sent the recorded tag ID, the received signal strength (RSS), and the time of detection to a central sensor station (Cellular Tracking Technologies, SensorStation, NJ, USA), which was uploaded to a cloud-based database for remote access. Tag localisations were then estimated using RSS-based multilateration based on previous calibrations of the relationship between RSS and distance in our study area (Tyson et al., 2024), which, when minimally three nodes detected a tag, produced estimated locations for each 15 second window (see Tyson et al., 2024, for more detail on the multilateration localisation method used in this study).

**Figure 2.**
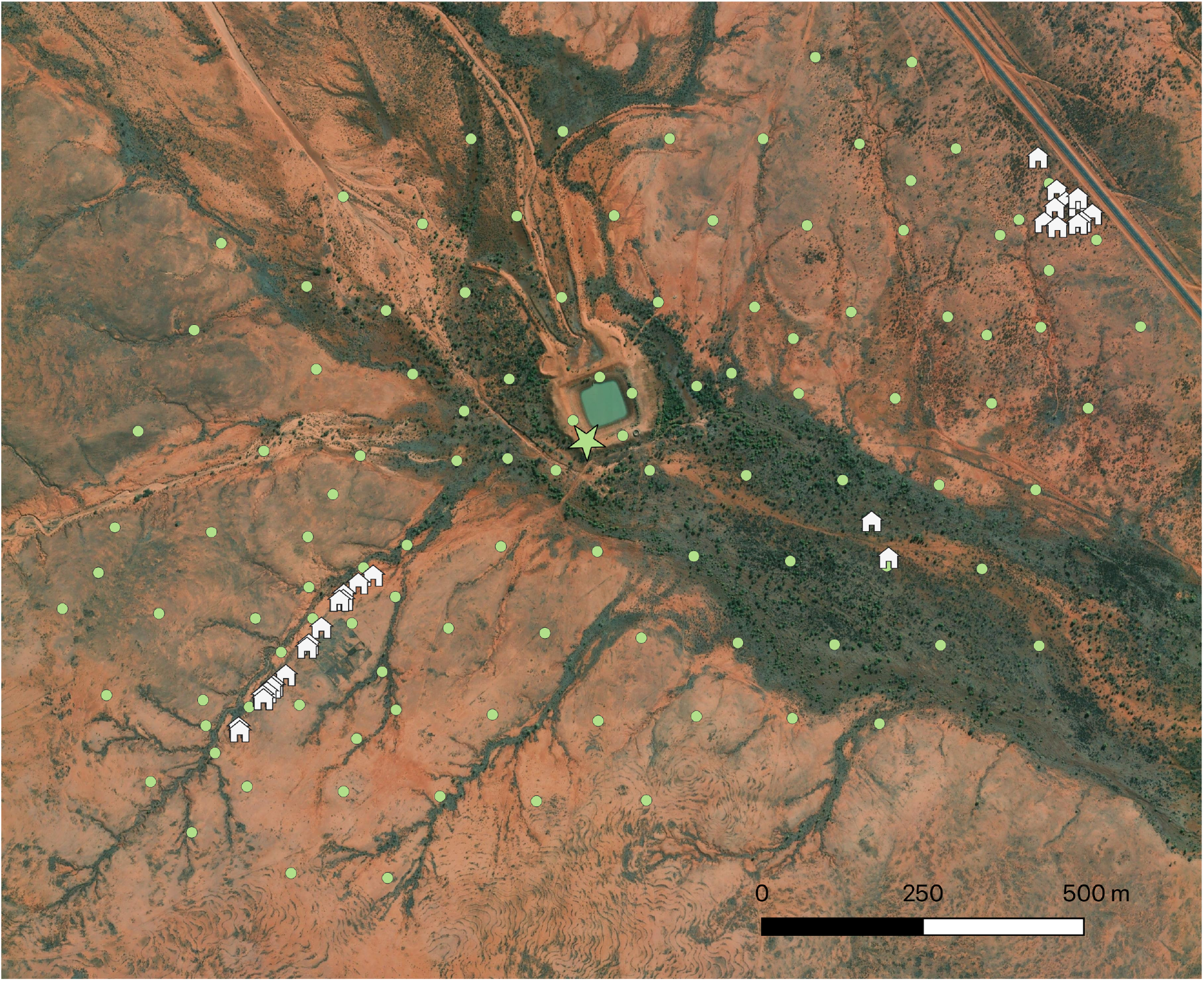
Tracking grid and nestbox setup in the study area ‘Gap Hills’ with nestboxes (white house icons) used by tracked families from three distinct breeding colonies spaced more than 500 m apart. Receiver nodes shown as green circles and the central sensor station as a green star at the central water dam. Basemap: Esri, World Imagery.

Tags were localised on average every 2 minutes (median = 0.5 mins, sd = 29 mins) during daylight hours. These smaller gaps were caused by birds not being picked up by at least three receiver nodes.

Larger gaps in the data were likely due to birds entering nestboxes or sitting in dense vegetation, resulting in insufficient solar power for the tag to signal, or due to birds temporarily leaving the grid. We tracked 161 birds (47 adult males, 41 adult females, 73 juveniles, of which 26 female, 40 male, 7 failed to be sexed) which included birds from 29 families, consisting of at least one parent and one juvenile that were localised simultaneously (see details below). Of these 29 families, we tracked both the mother and father for 18 families, only the father for 8 families and only the mother for 3 families.

### Data analysis

All data analysis was done in R (version 4.6.0, R Core Team, 2026). First, the tracking data was filtered to exclude periods after a tag’s antenna broke. A tag with a broken antenna is picked up less often by far fewer receivers, which results in fewer localisations with lower accuracy. In addition, since location estimates based on weak RSS values have lower accuracy, we further filtered the data by removing localisations that included a maximum RSS value of less than -80 dBm. This cutoff was chosen to balance the amount of data retained versus an adequate location accuracy (Speelman et al., 2025). Based on calibration data where we compared the estimated location to the ‘true’ position from GPS, the linear median error of these retained localisations was 23 m (range 1 – 45 m).

Next, we calculated the distance between individuals with simultaneous localisations to examine how juvenile distance to parents, siblings, and other colony members changed over juvenile age. As juvenile males start to practice song elements as early as day 25 (Roper and Zann, 2006), we decided to also include siblings and other juveniles, as they might influence each other’s song. To identify simultaneous fixes across individuals, we grouped localisations with the same timestamp using the function *group_times* in the package *spatsoc*. Dyads were filtered to include at least one juvenile. We identified the relationship within each dyad from the view of a focal juvenile to its dyadic partner, using a combination of nest and breeding area (Figure 2). This resulted in the dyadic relationships ‘sibling’ (two juveniles from the same nest), ‘parent’ (juvenile and adult from the same nest), ‘neighbouring peer’ (two juveniles from the same breeding colony, but from different nests), ‘neighbouring adult’ (juvenile and adult from the same breeding area, but different nests), ‘non-neighbour peer’ (two juveniles from different breeding colonies), and ‘non-neighbour adult’ (juvenile and adult from different breeding colonies). For each localisation per dyad, we added the age of the juvenile in days since hatching to evaluate changes in proximity over time. We calculated proximity as the distance in meters between simultaneous timestamps for every juvenile and every other tagged bird in the study area, using pairwise Euclidean distances through the function *edge_dist* in the *spatsoc* package. The sample size gradually declined over the tracking period, from 3,054 unique dyads at day 20 to 277 unique dyads at day 70, due to antenna breakage or tags stopping to signal at all as birds either left the tracking grid or got predated.

To determine how the proximity of juveniles to other birds changed over the course of the song learning period and whether this differed among relationships, we fitted a generalised additive mixed model (GAMM) using the function *bam* from the package *mgcv* (Wood, 2011). GAMMs are suited to model non-linear changes over a continuous predictor such as time, or age, in our case. We modelled the distance between juveniles and other birds as a function of juvenile age and dyadic relationship category, fitting a bam (big additive model), which is mgcv’s extension of gams for large datasets, using 3,363,484 observations (pairwise distance estimates). We specified a Gamma error distribution with a log link and estimated model parameters using fREML (fast Restricted maximum likelihood), and specified discrete=TRUE to speed up computation as this model is typically a sufficiently close approximation to the continuous form (Li and Wood, 2020). The relationship between distance and juvenile age was modelled using a separate smooth for each relationship category, with a basis dimension of k=20 for each smooth. The adequacy of basis dimensions was checked using *gam.check*. Relationship category was additionally added as a linear parametric term to allow different intercepts per category. Both individual ID and dyad ID were added as random effects to account for repeated observations of individuals, and repeated observations of the same dyad. Temporal autocorrelation was accounted for by specifying an AR1 correlation structure using the rho and AR1.start arguments in the function *bam*. Model diagnostics were checked using *gam.check*. In the same manner, we built two separate, smaller models to model juvenile-parent proximities depending on parent sex, as well as juvenile sex.

We then compared the estimated smooths between relationship categories using the *smooth_difference* function from the package *gratia* (Simpson, 2024), which estimates the difference between pairs of smooths across the range of juvenile age, allowing us to identify ages at which predicted distances differed between relationship categories. Differences were considered significant when the 95% confidence interval around the estimated difference did not include zero. From this, we identified periods during song learning when juveniles were significantly closer to, similarly distant from, or further from parents, siblings, and other birds in the colony. We used the same approach to assess whether juvenile sons and daughters differed in their distance to the parents, and whether juveniles differed in their distance to their mother versus their father across the period of song learning.

### Ethics

This project was approved by the Macquarie Animal Ethics Committee, and conducted under a New South Wales DPIE Scientific Licence SL100378, Ethics ARA 2018/027 and ABBBS Project 2697-01.

## Results

The average distance of juveniles to parents, siblings and neighbouring (i.e. colony member) peers and adults varied distinctly over the period of song learning (Fig. 3). At 20 days of age, right after fledging, a phase in which juveniles are mostly stationary while parents look for food, juveniles had a predicted average distance of 137 m (95% CI: 120-155 m) to their parents (Fig. 3, in red). Becoming more mobile with age, they gradually decreased the predicted average distance to their parents to 101 m (95% CI: 90-114 m) by day 35, then gradually increased the average distance to their parents to 283 m (95% CI: 193-372 m) at the end of the tracking period on day 70. Over the tracking period, juveniles and unrelated neighbouring adults or peers (other juveniles) gradually decreased their average distance, starting with 342 m (95% CI: 313-372 m) at day 20 to 153 m (95% CI: 132-174 m) at day 70 for juvenile--adult neighbour dyads (Fig. 3, in blue), and similarly, from 387 m (95%CI: 355-419 m) to 134 m (95% CI: 123-145 m) for juvenile-peer neighbour dyads (Fig. 3, in green). This pattern was similar when predicting the distances between juveniles to non-neighbouring adults and peers, although average distances were generally higher, with 669 m (95% CI: 613-725 m) at day 20 to 254 m (95% CI: 228-281 m) at day 70 and 699 m (95% CI: 642.755 m) at day 20 to 240 m (95% CI: 220-260 m) at day 70, respectively. Siblings stayed closest to each other immediately after fledging at a predicted distance of 43 m (95% CI: 38-48 m) at day 20, but over time the distance between siblings gradually increased up to day 50 after which it became more variable (Fig. 3, in yellow).

**Figure 3.**
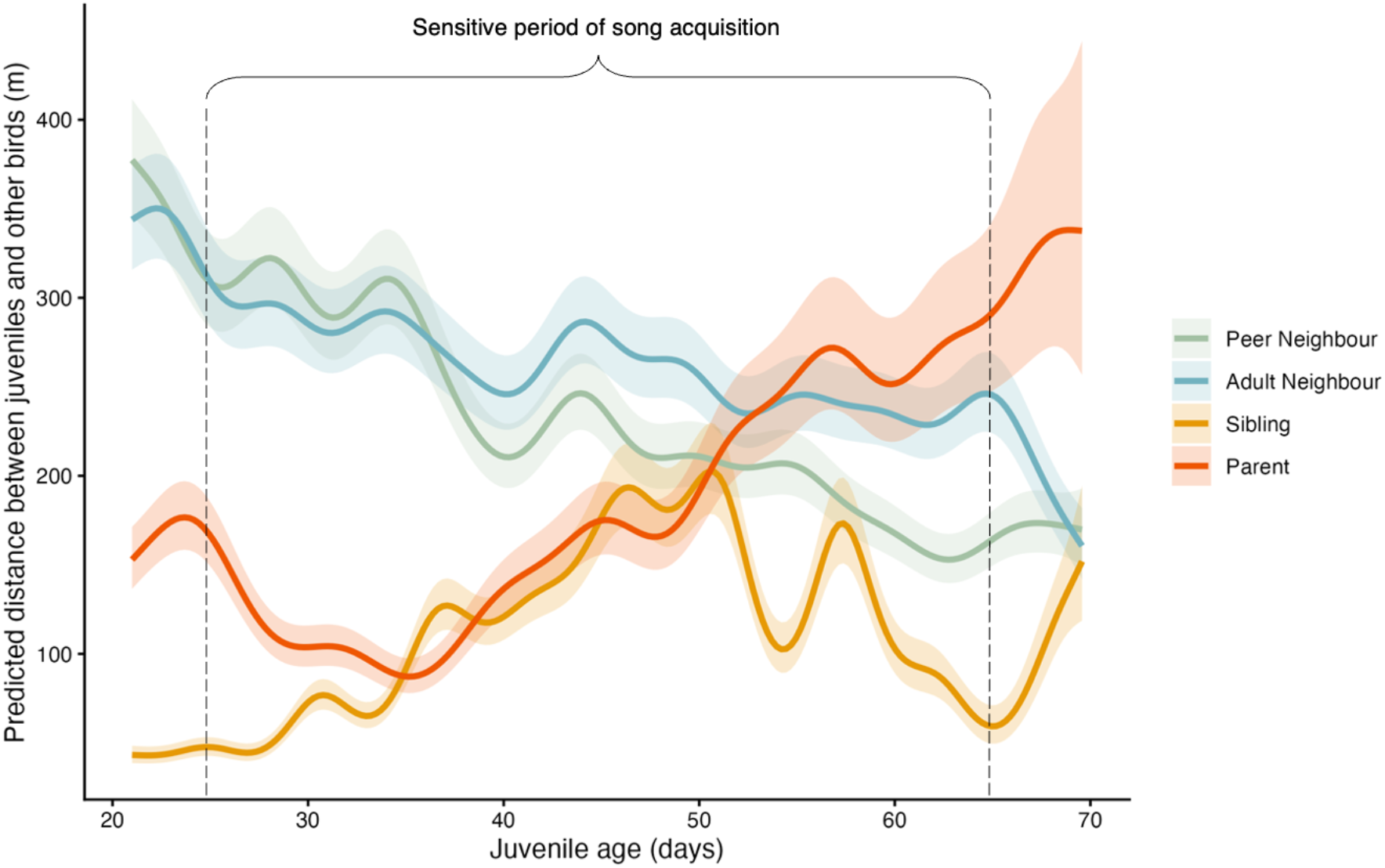
Predicted average distance between juvenile zebra finches and partners from different social categories as a function of juvenile age in days post-hatching. Lines show model-predicted average distances from a GAM with separate smooths for each relationship type (parents, siblings, peer neighbours and adult neighbours), shaded areas represent 95% confidence intervals around the predicted distances.

Comparing the juvenile-parent versus the juvenile-neighbour proximities, juveniles were significantly closer to their parents compared to adult neighbours until day 51, when, for the first time, their predicted distances no longer significantly differed as the confidence intervals cross 0 (Fig. 4A). This indicates that at 51 old, juveniles were as close to adult neighbours (other colony members) as they were to their parents. Similarly, juveniles were closer to their parents compared to peer neighbours until day 49, when they were not significantly closer to parents anymore. Juveniles remained similarly close to adult and peer neighbours for the first weeks after fledging but stayed slightly closer to peer neighbours over time (Fig. 3).

**Figure 4.**
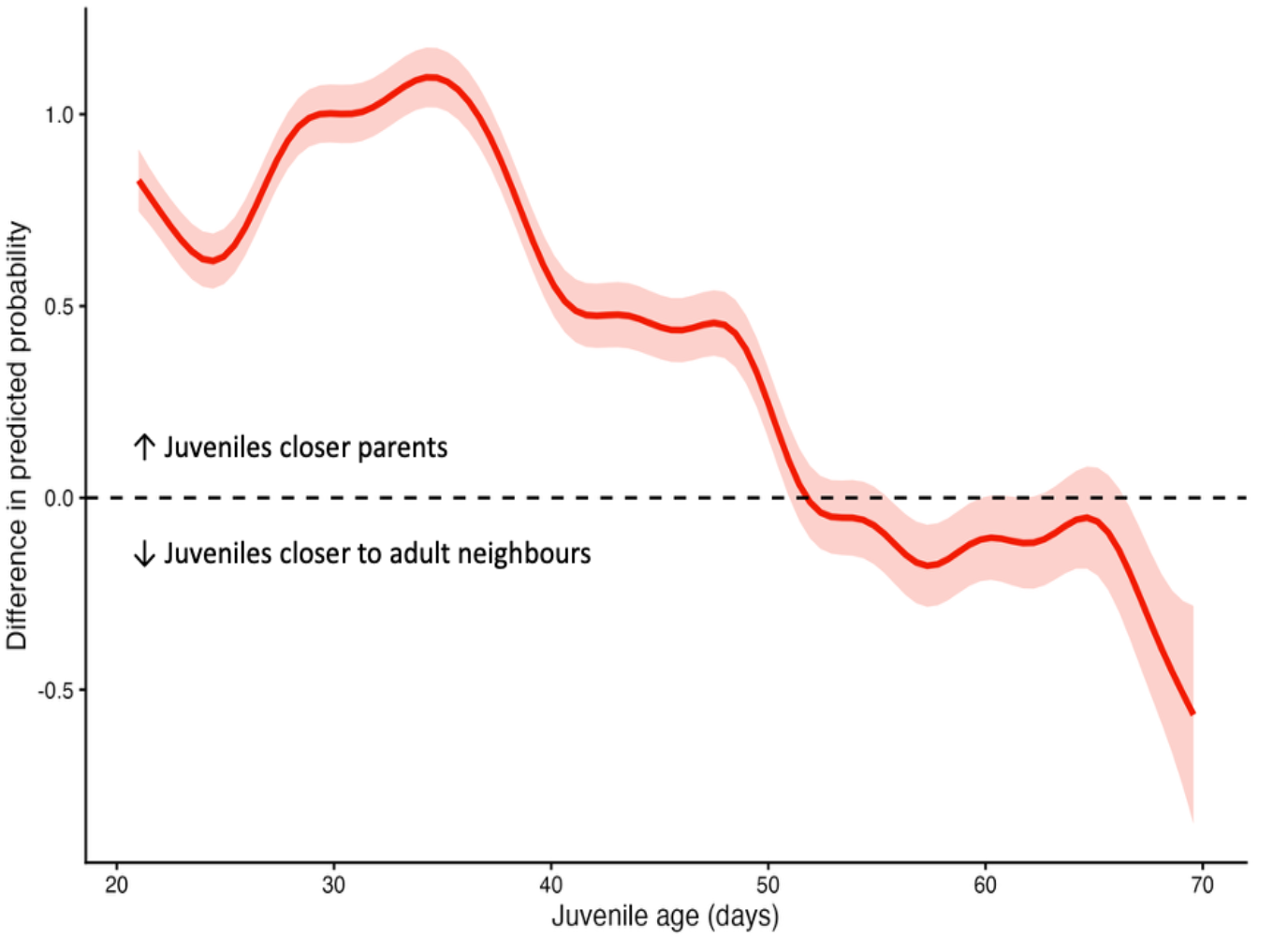
Juvenile distance to parents compared to juvenile distance to adult neighbours, confidence intervals overlapping 0 indicate no significant difference between smooths.

We detected no significant difference in proximities of juveniles to parents based on parent sex (Fig. 5A, 5B), indicating that juveniles stay as close to their mother as they do to their father over the sensitive period of song acquisition. Overall, male and female zebra finch juveniles did not significantly differ in how close they stayed to their parents across the period of song learning (Fig. 5C). Only briefly around day 40 and 50, female juveniles were significantly closer to their parents compared to male juveniles, though this difference was small (Fig. 5D).

**Figure 5.**
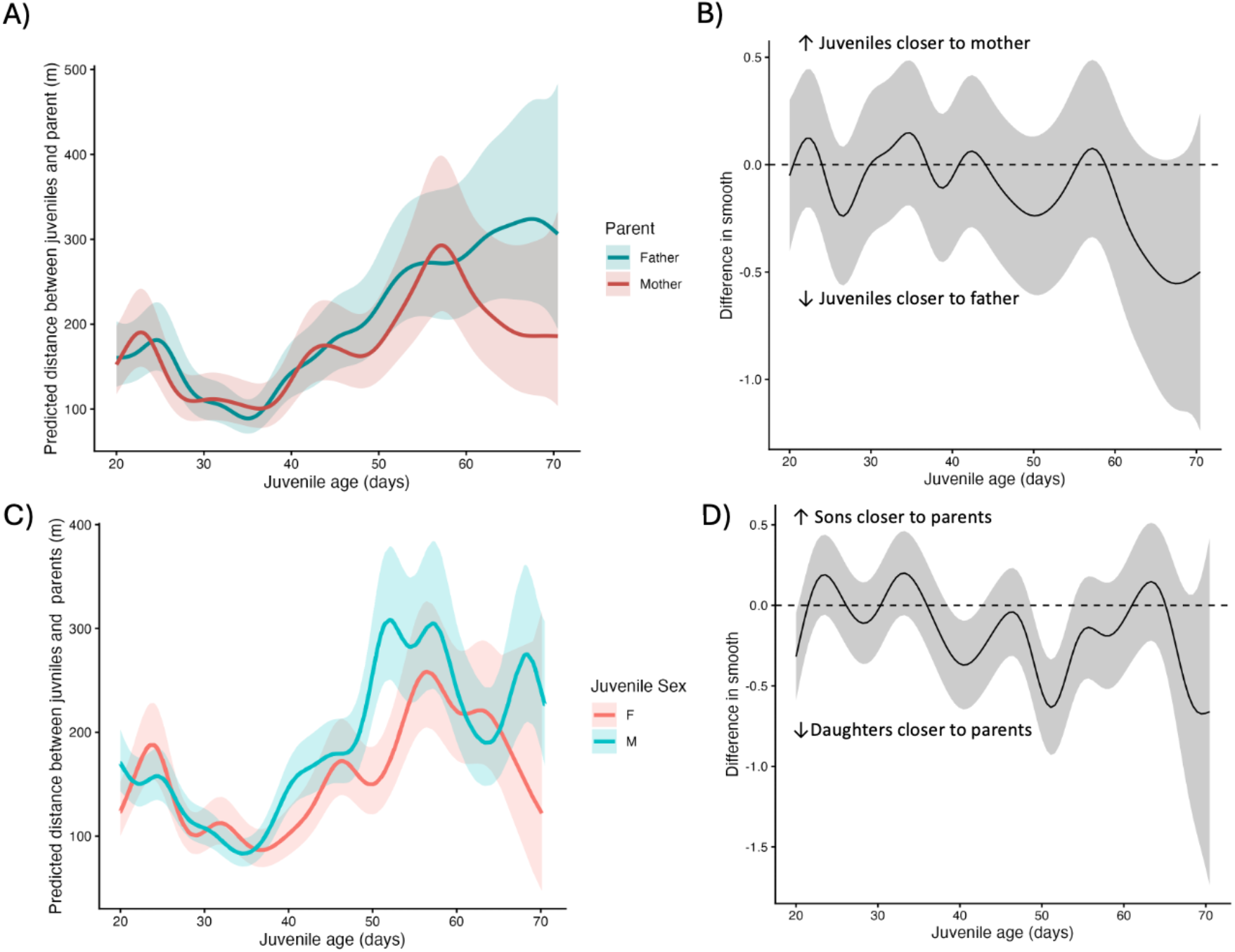
Predicted distance in meters of juveniles to parents depending on parent sex (A). Difference in smooths (B), confidence intervals overlapping 0 indicate no significant difference between mother and father distance to juvenile. Predicted distance in meters of juveniles to parents depending on juvenile sex (C). Difference in smooths (D), confidence intervals overlapping 0 indicate no significant difference between parents to juveniles based on sex, except for a brief window around age day 40 and 50.

## Discussion

Using automated radio-tracking of wild zebra finches during early development, we show how the key study model species for vocal learning is exposed to a more dynamic social environment than often assumed. Specifically, we show how proximity between juvenile zebra finches and their parents, siblings, and other colony members changed across the juvenile song learning period. We reveal a gradual decrease, followed by a gradual increase in distance between juveniles and parents, and most importantly, reveal that juveniles are as close to their parents as to other adult colony members while still well in the sensitive song learning period. Our findings thus suggest that juveniles are not exclusively exposed to the song of their father during the song learning period but rather experience the song of multiple potential tutors, identifying a mismatch between social conditions in many lab studies across the world compared to the natural conditions in the wild. As such, our findings thus provide a novel perspective on the social conditions during song learning.

Over the first two weeks after fledging, juveniles were in increasingly closer proximity to their parents. This is in line with our predictions, as newly fledged juveniles are largely sedentary, spending large parts of the day hidden in a bush with their siblings where they are provisioned by their parents (Zann, 1996). The observed gradual decrease in distance between juveniles and parents during the first two weeks post-fledging most likely reflects the juveniles’ increased mobility as they start to follow their parents around the landscape. At around day 35, juveniles reach nutritional independence from their parents (Zann, 1996). Our data support this, as from this age on, juvenile distance to parents gradually increased, suggesting that they began foraging on their own. This gradual separation from the parents is also theoretically expected, since while juveniles may benefit by being close to their parents for as long as possible, parents are expected to trade off investment in the current brood with future breeding efforts (Trivers, 1974). In zebra finches, such future breeding efforts may already start before day 35 of the current brood (Zann, 1996). From day 51, i.e. two weeks before the end of the sensitive period of song acquisition (day 65), juveniles were equally close to their parents as they were to other adults from the same breeding colony. This indicates that juveniles do not stay closely associated with their parents for the full duration of song learning and thus experience a strikingly different social context for song learning than studied in captivity. We found no prolonged significant difference in juvenile distance to parents depending on juvenile sex, suggesting that male and female offspring similarly benefit from being close to parents early in development, while forming closer associations with other birds later in the period of song learning. Juveniles were also similarly close to their mothers and fathers, suggesting that both parents are important social partners during early ontogeny. This is consistent with previous work showing strong spatial and temporal coordination between zebra finch partners, indicating that both parents are similarly involved in raising offspring (Mariette et al., 2012; Tyson et al., 2024).

Juveniles stayed closer to their siblings than to any other group, which suggests that the relationship between siblings may be especially relevant (Rebar et al., 2020). Staying close to siblings for longer throughout development might aid in inbreeding avoidance, as zebra finches in captivity have previously been found to only avoid inbreeding if birds can keep track of their kin (Ihle and Forstmeier, 2013). The prolonged close proximity between siblings observed in this study may also have implications for song learning. Juvenile males can influence each other through horizontal transmission of song (Gil et al., 2006; Honarmand et al., 2015; Tchernichovski and Nottebohm, 1998), and likewise female juveniles might be influenced for future mate choice or social preferences by their brothers’ song. As siblings stay in close proximity to each other for longer than they do to their parents, these effects might be stronger than previously assumed. In the laboratory, in broods with more male juveniles, songs are less similar to the father (Tchernichovski and Nottebohm, 1998) and element repertoire is smaller (Gil et al., 2006). Likewise, juveniles raised without exposure to the paternal song learned their song from a sibling (Derégnaucourt and Gahr, 2013). Juvenile songs even showed greater similarity to each other than to the song of the father that served as the model, but that was physically separated from the juveniles (Honarmand et al., 2015). Furthermore, different song characteristics were affected differently by the social conditions, such as peer group size and sex ratio, during song learning (Honarmand et al., 2015). For future studies, it would thus be valuable to compare a juvenile’s song to their sibling’s song in comparison to the father’s song as well as to other males that they are exposed to during song learning. While both siblings might initially copy elements from their father, and later from a range of tutors, they might further influence each other’s songs.

Throughout the period of song learning, we found a gradual decrease in distance between juvenile zebra finches and other juveniles and adults in the colony. This suggests that as juveniles age, they become closer to other birds that are not their immediate family. This shift, together with being progressively farther away from parents throughout the sensitive period for song learning, shows that juveniles are exposed to other individuals who could potentially serve as song tutors. Yet, zebra finch juveniles are likely not learning song from just any singing adult around them, instead, they learn from tutors who direct song towards them (Chen et al., 2016), or directed song towards a female (Roper and Zann, 2006). Songs directed at tutees differ from songs sung in other contexts, for example, by a change in the temporal song structure, which in turn increases the attention of juveniles to the tutor (Chen et al., 2016). Therefore, it might not only matter which other individuals juveniles are close to, but to whom they give their attention. Juveniles in the laboratory preferentially learn from tutors with whom they have most social interactions, regardless of kinship (Mann and Slater, 1995; Williams, 1990). To better understand such social learning strategies (Kendal et al., 2018) studying proximity within a family and among neighbours during the song learning period can be complemented with direct, visual observations at social hotspots in addition to comparing audio recordings from adults and male juveniles.

Overall, our findings on the social environment of juveniles and how it changes during song learning help to contextualise studies from controlled laboratory conditions relative to the natural environment of zebra finches. Song learning in zebra finches is a widely used and impactful model across many research fields, such as linguistics, neurobiology, behaviour, and evolution (Hauber et al., 2021), where individuals are typically exposed to a very limited number of tutors. Consequently, our findings that during the song learning period, in nature, juveniles are naturally exposed to many tutors have broader implications as to how to interpret the functional implications of the song learning process in a highly controlled laboratory environment. Selection on song learning in males may favour not copying the father’s song, while in females, it may favour not developing sexual and social preferences for their father’s song, as is generally suggested by laboratory studies (Riebel, 2009). Instead, the mechanisms of song learning and song preference, including the duration and timing of the most sensitive period, might unfold very differently under natural conditions, where juveniles encounter multiple tutors and develop diverse social connections. Such exposure to multiple tutors likely also facilitates the striking individual distinctiveness of zebra finch song (Böhner, 1983; Woodgate et al., 2012; Zann, 1990), which in turn may be crucial for partners to keep track of each other in their fission-fusion social system (Hagedoorn et al., 2026). Overall, our study on zebra finches has wide implications for interpreting many of the laboratory findings in a functional framework and encourages including more socially complex contexts in the lab to be able to identify if and how song learning mechanisms are affected by the social conditions during this period. More broadly, our study also has wider implications for studies on other species in captive conditions as it highlights the value of incorporating more natural social settings into laboratory studies to ensure ecologically relevant and generalizable insights (Webster and Rutz, 2020). Studying behavioural processes under more natural social conditions will allow us to better understand and integrate the biological mechanisms and conditions under which particular behaviours have evolved and function.

## Acknowledgements

We thank Katja Hagedoorn, Marie-Anne Bellon and Tis Voortman for their help with field data collection. Thanks to Garry Dowling and Vicki Dowling for their logistical support in the field.

